# Cryo-EM structure of an early precursor of large ribosomal subunit reveals a half assembled intermediate

**DOI:** 10.1101/242446

**Authors:** Dejian Zhou, Xing Zhu, Sanduo Zheng, Dan Tan, Meng-Qiu Dong, Keqiong Ye

**Author notes:** To whom correspondence should be addressed., Tele: 86-10-64887672. These authors contribute equally to this work (Dejian Zhou, Xing Zhu, Sanduo Zheng).

## Abstract

Assembly of eukaryotic ribosome is a complicated and dynamic process that involves a series of intermediates. How the highly intertwined structure of 60S large ribosomal subunits is established is unknown. Here, we report the structure of an early nucleolar pre-60S ribosome determined by cryo-electron microscopy at 3.7 Å resolution, revealing a half assembled subunit. Domains I, II and VI of 25S/5.8S rRNA tightly pack into a native-like substructure, but domains III, IV and V are not assembled. The structure contains 12 assembly factors and 19 ribosomal proteins, many of which are required for early processing of large subunit rRNA. The Brx1-Ebp2 complex would interfere with the assembly of domains IV and V. Rpf1, Mak16, Nsa1 and Rrp1 form a cluster that consolidates the joining of domains I and II. Our structure reveals a key intermediate on the path to the establishment of the global architecture of 60S subunits.

## Introduction

Correct formation of the ribosome is essential for protein synthesis in cells. The ribosome is composed of a small 40S and large 60S subunit (SSU and LSU) in eukaryotes. Ribosome assembly is a highly complicated process that engages more than 200 protein assembly factors (AFs) and many snoRNAs (1). These factors temporarily associate with ribosomal subunits and drive their maturation. In the yeast *S. cerevisiae*, assembly of both subunits begins in the nucleolus with the transcription of a long precursor rRNA (pre-rRNA) that encodes 18S, 5.8S and 25S rRNAs and four external and internal transcribed spaces (ETS and ITS). The 5’ region of the pre-rRNA is co-transcriptionally assembled into the 90S pre-ribosome or SSU processome, which is the early precursor of SSU (2–6). Following cleavage of the pre-rRNA at A0, A1 and A2 sites, the 90S is released from the pre-rRNA and transformed into a pre-40S ribosome. The cryo-EM structures of 90S pre-ribosome show that the nascent 40S subunit is assembled into several isolated units that have yet to acquire the global architecture of 40S subunit (7–11).

The 60S subunit is composed of 25S, 5.8S and 5S rRNAs and 46 ribosomal proteins (RPLs). The 25S/5.8S rRNAs are divided into six domains that intertwine into a monolithic structure (12, 13). The earliest pre-60S ribosome is co-transcriptionally formed on the 3’ LSU region of pre-rRNA in a stepwise manner (14). The pre-60S progressively maturates as it transits from the nucleolus to nucleoplasm and cytoplasm. The maturation of pre-60S is coupled to dynamic association of approximately 90 assembly factors, hierarchical incorporation of RPLs and sequential processing of the ITS1 and ITS2. The development of pre-60S at the nucleoplasm and cytoplasm has been characterized to great extents, particularly from recent cryo-EM studies (15). An early nucleoplasmic state is characterized by a foot structure assembled on the partially processed ITS2 and an immature central protuberance (CP) (16–18). Further evolvement of pre-60S leads to removal of the foot, maturation of the CP and association of the Rix1-Ipi1-Ipi2 subcomplex and the AAA-ATPase Rea1(19). Rea1 hydrolyzes ATP and drives the release of Ytm1 and Rsa4 (20, 21). The pre-60S acquires competence for nuclear export at the late nucleoplasmic stage and undergoes final maturation in the cytoplasm (22–26).

In all the determined cryo-EM structures of pre-60S, the structural core already takes the shape of mature subunit. Therefore, the key question how is the global architecture of LSU is established is still unknown. In this study, we determine the cryo-EM structure of pre-60S at an early nucleolar state. The structure is only half assembled at domains I, II and VI and represents a key intermediate in the formation of global architecture of LSU.

## Results

### Structure determination

We affinity purified pre-60S via the bait protein Rpf1 fused with a C-terminal tandem affinity purification (TAP) tag. Mass spectrometry analysis show that the Rpf1-TAP particle displayed a similar protein composition as the previously characterized nucleolar Ssf1-TAP and Nsa1-TAP particles (14, 27, 28) (Figure S1, Supplementary Dataset 1). Cryo-EM images were collected in a Titan Krios 300 kV electron microscopy equipped with a direct electron detector (Figure S2). After three rounds of 3D classification, 98155 particles were selected to reconstruct a density map at an overall resolution of 3.73 Å (Figure 1A and Figure S3). Large side chains of protein and RNA bases are resolved in the structure core, whereas some peripheral regions including many AFs are of lower resolution (Figure S4). The structure was built based on the cryo-EM structure of Nog1-TAP pre-60S (18), crystal structures (Brx1, Ebp2, Nsa1), homology models (Has1 and Rpf1) and de novo modeling (Mak16 and Rrp1) (Figure S4, Table S1). The Rpf1-TAP sample was also analyzed by chemical crosslinking and mass spectrometry (CXMS), yielding 31 intermolecular and 107 intermolecular crosslinks (Supplementary Dataset 2). These crosslinks assisted the assignment of AFs (Table S1). The final model was refined in real space and displayed good geometry (Table S2).

**Figure 1.**
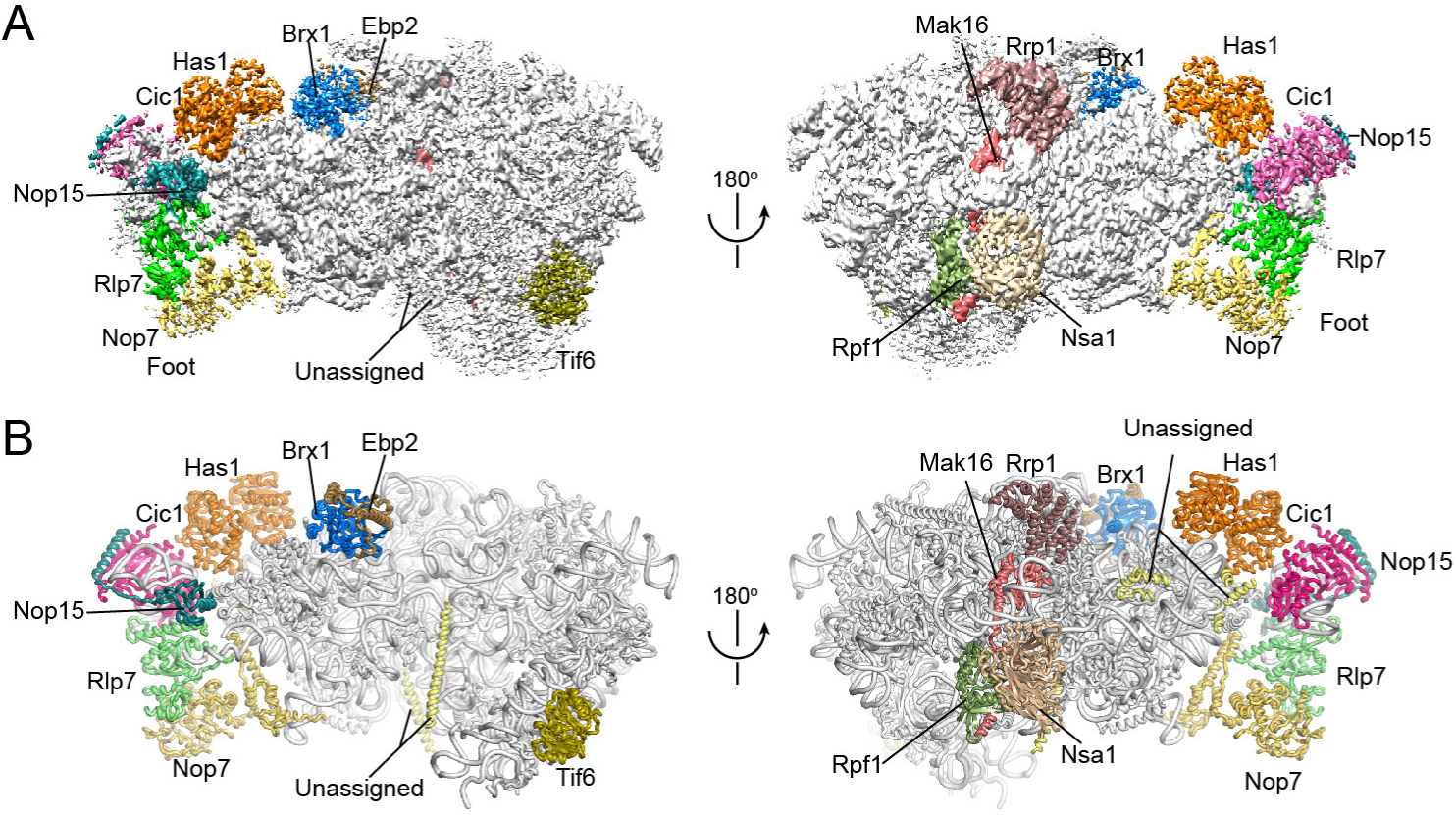
Cryo-EM structure of Rpf1-TAP pre-60S. (A) Cryo-EM density map in two opposite views. The densities for AFs are colored coded. (B) Structural model in the same views as A. The rRNAs and RPLs are colored silver and the AFs are color coded.

### Overall structure

The cryo-EM density map and structural model of Rpf1-TAP pre-60S are shown in Figure 1. The most striking feature of the structure is that only half of LSU is present. Domains I, II and VI assemble into a native-like substructure, but domains III, IV and V are totally absent (Figure 2). A few peripheral elements in domain II that form the CP (H38, ES12) and the P stalk (H43, H44) and interact with domains III, VI and V (H33-H35) are also disordered (Figure 2E). Therefore, the structure represents an early assembly intermediate before the global architecture of 60S is established. A total of 12 AFs and 19 RPLs were modeled in the map (Figure 1B and 2C-D). Five AFs Nop7, Rlp7, Nop15, Cic1 and Tif6 are also present in the early nucleoplasmic pre-60S (16–18) and adopt similar conformation in the Rpf1-TAP pre-60S. Seven AFs Has1, Brx1, Ebp2, Rrp1, Mak16, Nsa1 and Rpf1 were newly located (Figure S3).

**Figure 2.**
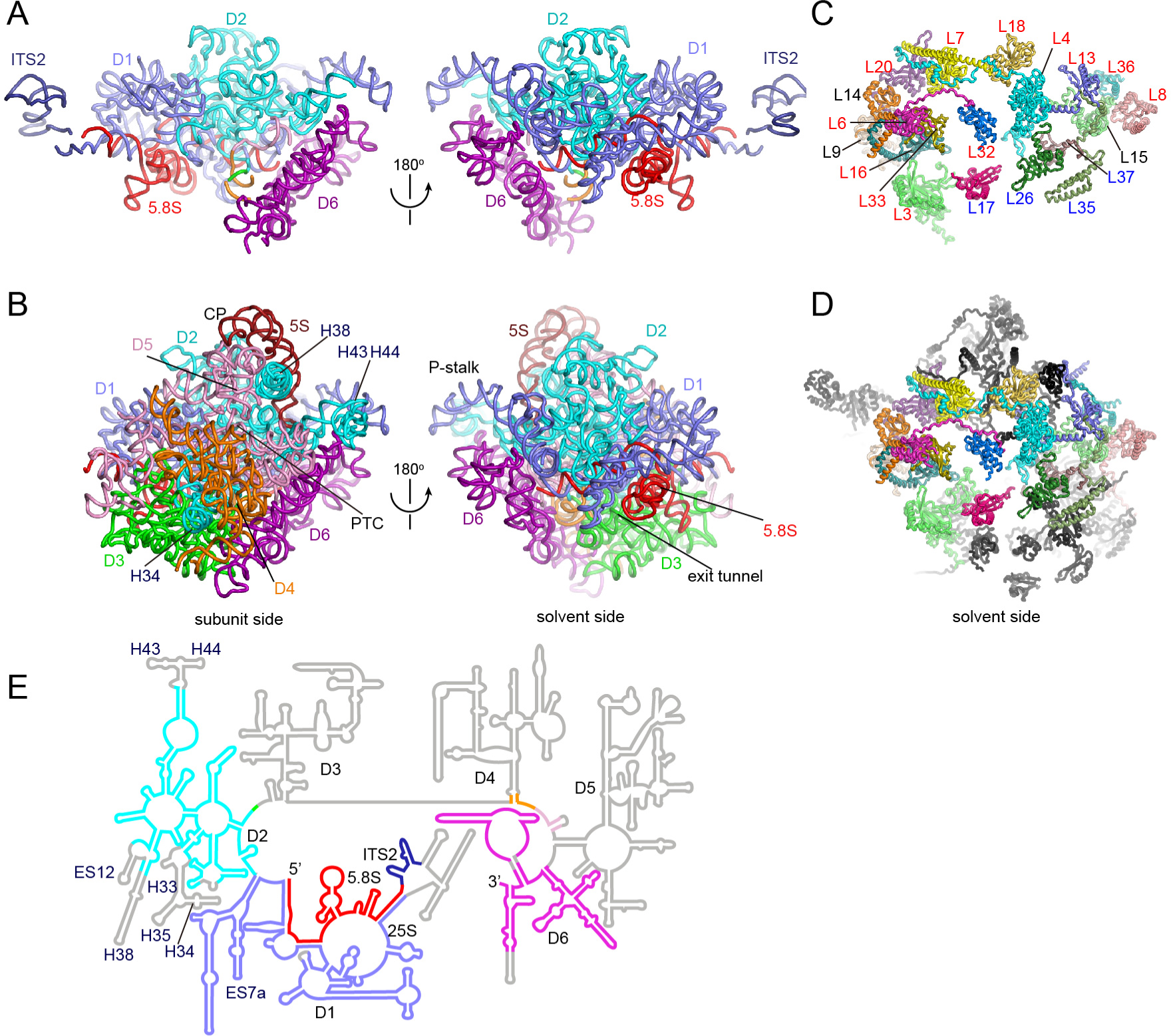
Structure of rRNAs and RPLs in Rpf1-TAP pre-60S and mature 60S subunit. (A-B) Structure of rRNAs in Rpf1-TAP pre-60S (A) and mature 60S (B). The subunit and solvent side views are shown. Domains I-VI of 25S are colored slate, cyan, green, orange, pink and purple. ITS2, 5.8S and 5S are colored deepblue, red and brown. Central protuberance (CP), peptidyl transferase center (PTC), exit tunnel and P-stalk are labeled. (C-D) RPLs in Rpf1-TAP pre-60S (C) and mature 60S (D). The solvent side view is shown. RPLs present in pre-60S are color coded and those missing in pre-60S are shown in black in D. The early, middle and unclassified RPLs are labelled in red, blue and black. (E) Secondary structure model of 5.8S, ITS2 and 25S RNA. The modeled regions are colored as in A-B and the unmodeled regions are colored grey.

### Assembly of ITS2

The N-terminal region of ITS2 is associated with Cic1, Nop15, Rlp7 and Nop7, forming the foot structure (Figure 1 and Figure 3A). The foot structure is similar with that in the nucleoplasmic pre-60S structures (16–18), except that Nop53 is not present (Figure 3B). Nop53 recruits the exosome for processing 7S pre-rRNA (29, 30). Nop53 is of low abundance in Rpf1-TAP particle (Figure S1) and should associate at a later stage.

**Figure 3.**
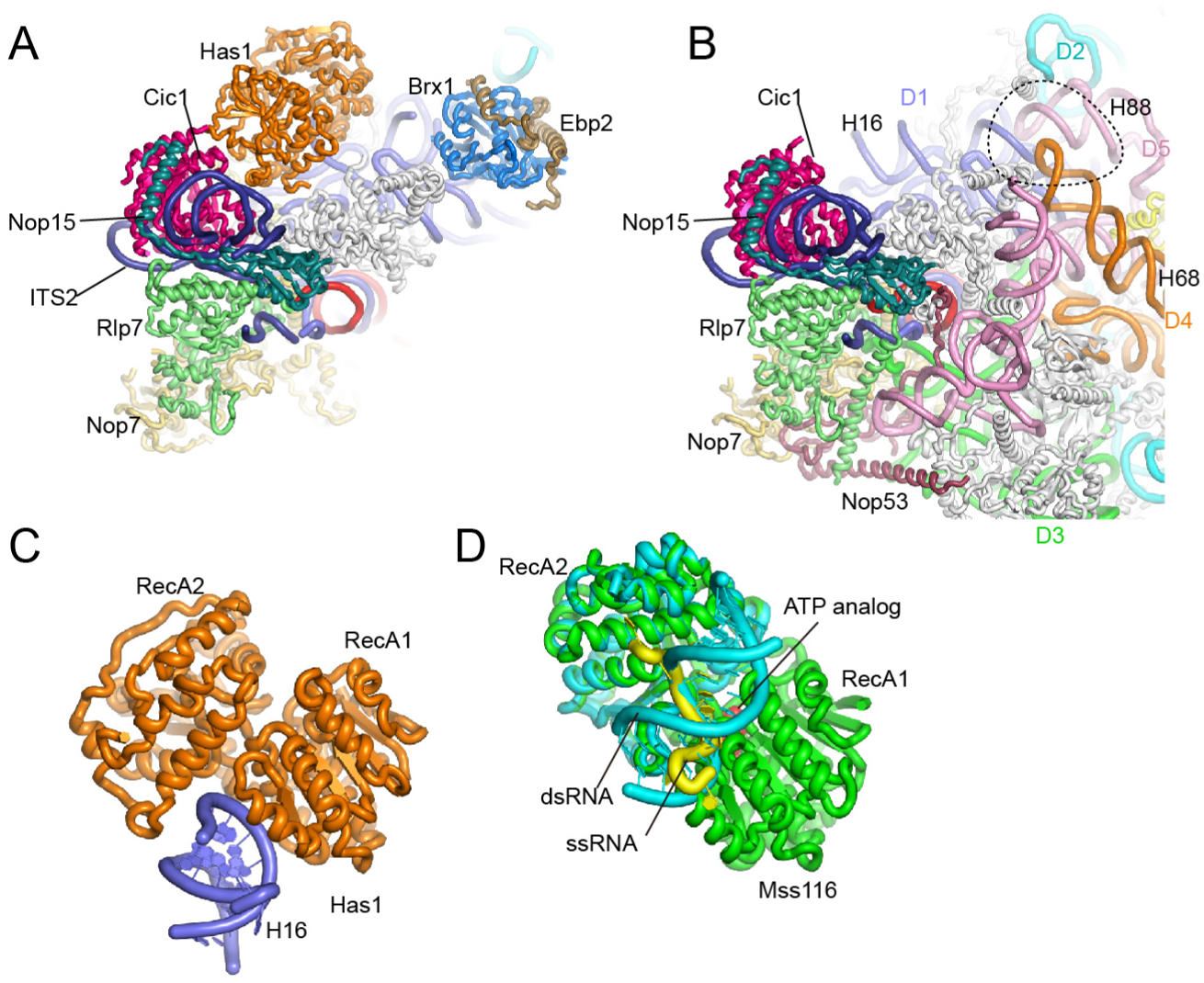
AFs bound to ITS2 and domain I. (A) Structure of Rpf1-TAP pre-60S near the ITS2. AFs, ITS2, 5.8S and individual domains of 25S rRNAs are color coded. All RPLs are shown in silver. (B) Structure of the Nog1-TAP pre-60S (PDB code 3JCT) shown in the same view as A. The dashed circle indicates the binding site of Brx1 and Ebp2 that would clash with H68 and H88. (C) Structure of Has1 bound to H16. (D) Superimposed structures of Mss116 in complex with ssRNA and dsRNA. Mss116 is aligned to Has1 in C by RecA1 domain. The ternary complex structure (PDB code 3I5X) is composed Mss116 (green), an ssRNA (yellow) and an ATP (red). The binary complex containing the RecA2 domain of Mss116 bound to an RNA duplex (PDB code 4DB2) is colored cyan and aligned to the ternary complex.

### Has1

Has1 is a DEAD-box RNA helicase involved in both small and large ribosomal subunit assembly (31, 32). RNA helicases are thought to unwind RNA structures or disassemble RNA-protein complexes, hence driving the evolvement of pre-ribosomes. In the cryo-EM structure, Has1 is situated next to Cic1 and rides on H16 of 25S rRNA with its two RecA domains (Figure 3A). DEAD-box helicases unwind short RNA duplexes by local strand separation (33). In a proposed mechanistic model (34), RecA2 first associates with the RNA duplex and binding of ATP to RecA1 then induces closure of two RecA domains, causing RecA1 to displace one RNA strand. In the pre-60S structure, H16 is bound to Has1 at a site different from where single- and double-stranded RNAs are bound to DEAD-box helicases (Figure 3C-D) (34, 35). In addition, the ATP-binding pocket of Has1 is empty and its two RecA domains are not arranged in the same closed conformation as in ternary complex structures of helicase, ssRNA and ATP analog. These structural features suggest that H16 serves as a docking site, rather than an unwinding substrate, for Has1.

### Structure of the Brx1 and Ebp2 complex

Brx1 is a Brix domain protein and interacts with Ebp2 (36, 37). We determined a crystal structure of Brx1 (residues 26–255) in complex with Ebp2 (residues 195–293) at 2.3 Å resolution with de novo phasing (Table S3). Brx1 adopts an α/β fold with pseudo 2-fold symmetry (Figure 4A), like other Brix domain proteins (38–42). The structural core of Brx1 is composed of two curved β-sheets that from a half-open β-barrel sandwiching two α-helices (α1 and α3). The surface of the N-terminal sheet holds helix α2, whereas the surface of the C-terminal sheet makes an intermolecular interaction with helix α2’ of Ebp2. In addition, the Ebp2 sequences flanking helix α2’ wrap around the Brx1 structure, forming an extensive interface. The α2’ helix of Ebp1 packs against the N-terminal sheet of Brx1 and the β1’ strand of Ebp2 expands the C-terminal sheet of Brx1.

**Figure 4.**
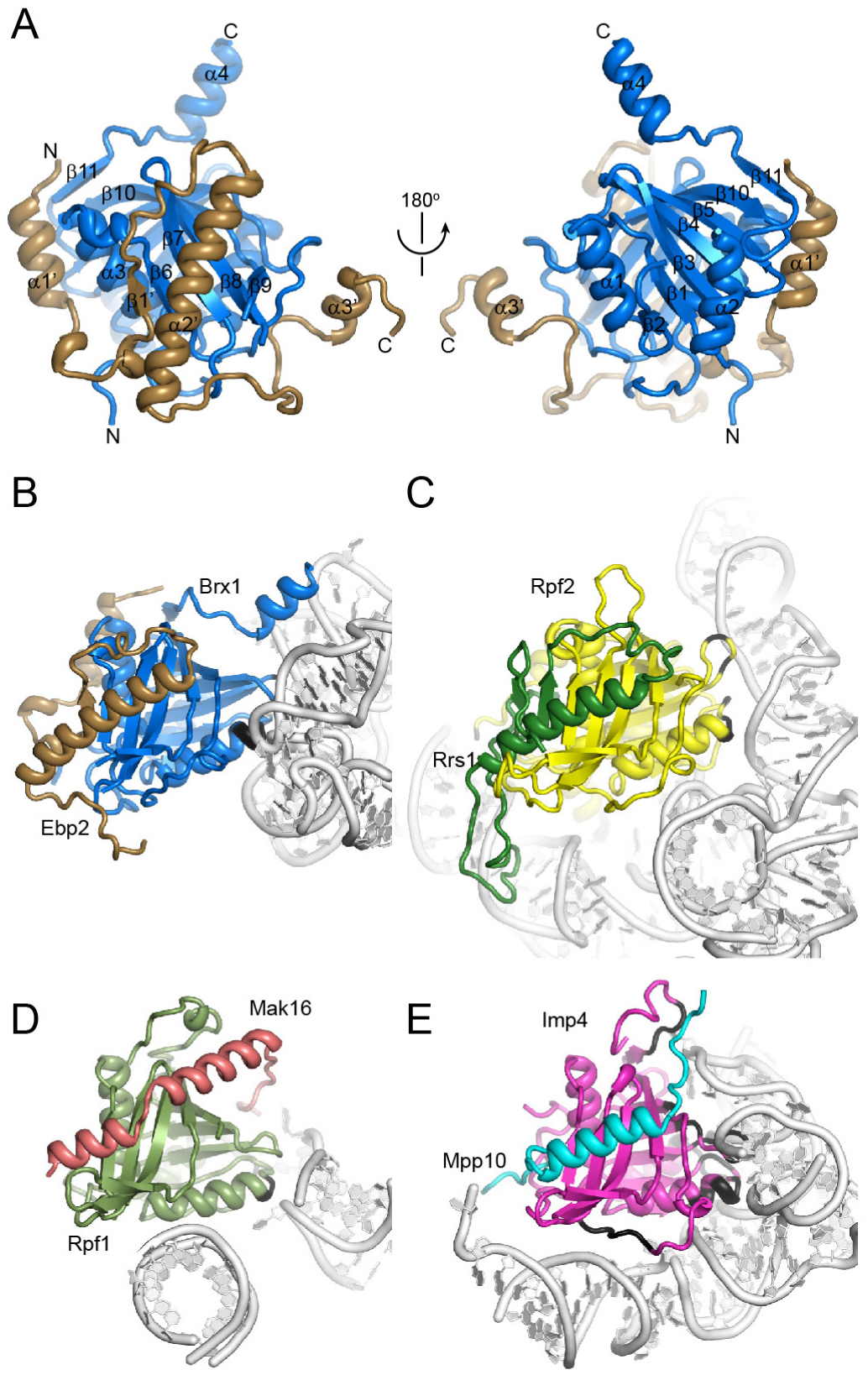
Structure of the Brx1-Ebp2 complex. (A) Crystal structure of the Brx1-Ebp2 complex shown in two opposite views. The N- and C-termini and secondary structures are labeled. (B-D) Comparison of protein and RNA interactions of four Brix domain proteins in the context of pre-ribosome structures. Proteins are color coded and RNAs are colored in silver. The protein atoms within 5 Å of any RNA atom are colored black. The Brx1-Ebp2 complex in the Rpf1-TAP pre-60S (B). The Rpf2-Rrs1 complex in the Nog1-TAP pre-60S (PDB code 3JCT) (C). The Rpf1-Mak16 complex in the Rpf1-TAP pre-60S (D). The Imp4-Mpp10 complex in the 90S pre-ribosome (PDB code 5WLC) (E).

In the cryo-EM map, the Brx1-Ebp2 complex perfectly fits into a density bound to domain I (Figure 1 and S4B). Brx1 interacts with a flat surface formed by H13 and H21. The position of Brx1 would clash with H68 and H88 in domains IV and V in the fully assembled 60S (Figure 3B). Thus, the Brx1-Ebp2 complex must be released from the current position for ribosome assembly to proceed.

There are six Brix domain proteins Imp4, Rpf2, Rpf1, Brx1, Ssf1 and Ssf2 in the yeast S. cerevisiae. Imp4 is involved in small subunit assembly, whereas the others function in large subunit assembly. The structures of Rpf2, Imp4, Rpf1 (see below) and Brx1 have now been determined in the context of pre-ribosomes (7–11, 16, 18), allowing for comparison of their conserved and variable features in protein and RNA recognition (Figure 4B-E). The four proteins all bind an α-helix of their partner proteins with the C-terminal sheet of Brix domain. Outside the core interface, the intermolecular interaction is highly variable. Brx1 and Rpf2 form an extensive interface with their binding partners Ebp2 and Rsr1, but Rpf1 and Imp4 primarily bind only a single α-helix of Mak16 and Mpp10. The RNA binding interface is also extremely diverse for the four Brix proteins. No consensus sequence or structure is apparent for the bound RNAs. Nevertheless, one protein region that consistently binds RNA is located at the junction between strand β1 and helix α1 and the turn connecting strands β3 and β4. The diverse interactions with protein and RNA allow each Brix protein play specific roles in ribosome assembly.

### D2 factors

Rrp1, Mak16, Rpf1 and Nsa1 form a cluster bound to a junction region of domains I and II at the solvent side (Figure 5A-F). The four proteins are co-assembled to a pre-rRNA fragment ending at domain II (termed as D2 factors) (14), consistent with their binding sites. These AFs are not present in the nucleoplasmic pre-60S and Nsa1 is released by the AAA ATPase Rix7 (28).

**Figure 5.**
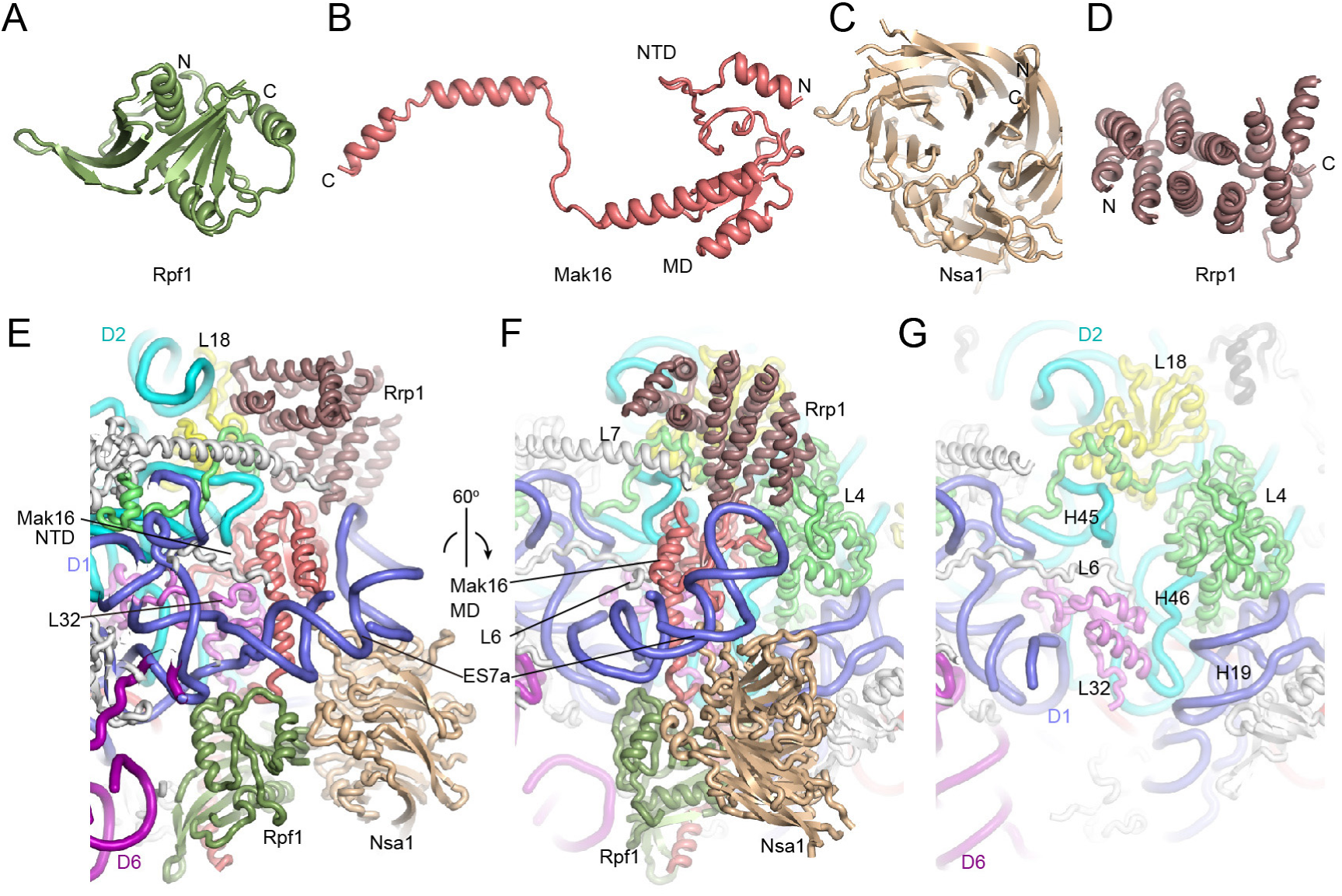
Assembly of D2 factors. (A-D) Structure of Rpf1 (A), Mak16 (B), Nsa1 (C) and Rrp1 (D). The N- and C-termini are labeled for each protein. The N-terminal domain (NTD) and middle domain (MD) of Mak16 are indicated. (E-F) Two views of Rpf1, Mak16, Nsa1 and Rrp1 assembled at the junction of domains I and II. AFs and individual domains of 25S rRNAs are color coded. L4, L18 and L32 are colored green, violet and yellow and other RPLs are shown in silver. (G) Structure of mature 60S subunit (PDB code 4V88) in the same view as F.

Mak16 occupies a central position of the cluster and is mostly buried. Mak16 is composed of a knot-like N-terminal domain (NTD), a two-layered middle domain (MD) and an extended C-terminal helix (Figure 5B). The NTD fits snuggly into a cavity formed by H45 and H46 and ribosomal proteins L4 and L32. The MD is covered by the first helix of expansion segment 7 (ES7a) RNA. The C-terminal α-helix of Mak16 binds the Brix domain protein Rpf1. This interaction was also demonstrated by yeast 2-hybrid, in vitro pull-down and gel filtration assays (Figure S5) (43, 44). Besides binding Mak19, Rpf1 also binds L32, Nsa1 and the base of helix ES7a. Nsa1 contains a WD domain (45) and is docked at H19, ES7a and Rpf1. Rrp1 folds into a super-helical structure and contacts L4, L18, the N-terminus of L7, Mak16 and RNA helix ES7a. By binding both domains I and II, these D2 factors appear to strengthen the joining of the two domains at early assembly stages when many interactions in the mature subunit are missing.

The ribosomal structure at the D2 factor binding region closely resembles the mature subunit (Figure 5F). However, in the mature subunit, the N-terminal tail of L6 inserts into the cavity initially occupied by Mak16 and ES7a becomes disordered in the absence of AF contacts. ES7a is a eukaryote-specific element in 25S rRNA and appears to function in ribosome assembly, which differs dramatically between bacteria and eukaryotes, rather than in protein translation.

## Discussion

We have determined the cryo-EM structure of pre-60S at an early nucleolar state. The structure demonstrates that the global architecture of 60S is first established by domains I, II and VI that constitute the major part of the solvent shell of 60S. Part of the solvent shell (domain III) and the entire inner layer at the subunit interface (domains IV and V) have not been integrated. The functional sites of 60S, peptidyl transferase center and polypeptide exit tunnel, are not assembled at this stage. The absence of domains III, IV and V does not mean that these domains are completely unfolded. They could fold and associate with AFs and RPLs (14), but are too flexible to be visualized in the cryo-EM structure. The Rpf1-TAP particle contains ~40 AFs of appreciable abundance (Figure S1). Those AFs not present in our structure may be flexible or associate with other states of pre-60S. The map also contains unassigned structures and unmodeled weak densities.

In the Rpf1-TAP pre-60S, domains I, II and VI together with 19 RPLs are tightly packed as in the mature subunit. All AFs barely interfere on the formed ribosomal structure. By contrast, in the 90S pre-ribosome, all four domains of 40S subunit are isolated and organized by extensive interactions with a large number of AFs, the 5’ ETS and U3 snoRNA (7–11).

Many AFs and RPLs have been classified according to their role in pre-rRNA processing (1, 46–49). A3-factors (or early factors) are required for processing of 27SA3 to 27SB pre-rRNA, B-factors (or middle factors) are required for C2 cleavage of ITS2 and late factors are required for processing of 7S pre-rRNA. Base on functional classification, binding dependence and structural location of RPLs, a hierarchical model has been proposed for 60S assembly with assembly beginning at the solvent side (49). The Rpf1-TAP pre-60S structure is half assembled at the solvent side and contains 8 of 13 AFs and all of 12 RPLs that are classified as early A3-factors (Figure 2C and Figure S1). This suggests that the determined structure corresponds to the predicted early assembly intermediate and is required for ITS1 processing.

By analyzing the association of AFs and RPLs to plasmid-derived pre-rRNA fragments of increasing length, the earliest precursor of 60S was found to form in a stepwise manner (14). The D2 factors Rpf1, Mak16, Nsa1 and Rrp1, which are initially recruited by a pre-rRNA fragment ending at domain II, are found to bind at the junction of domains I and II. This suggests that domains I and II could already associate in the absence of other domains. This leads to a sequential assembly model for the Rpf1-TAP pre-60S. Domains I and II first combine into a structural core that is late joined by domain VI.

## Materials and methods

### Purification of Rpf1-TAP particles

The Rpf1-TAP strain (BY4741, RPF1-TAP::HisMX3) was purchased from Open Biosystems. Six liters of cells were grown at 30 °C in YPD medium (1% yeast extract, 2% peptone, 2% glucose) to OD_600_ of 1–2. The cells were harvested and resuspended in lysis buffer (20 mM HEPES-K pH 8.0, 110 mM KOAc, 40 mM NaCl, 0.1% NP-40) supplemented with one tablet of EDTA-free protease inhibitor cocktail (Roche). The cells were lysed by a high pressure cell disruptor (JNBIO) and clarified by centrifugation at 6,000g for 10 min. The supernatant was filtered through a 0.45 µm membrane and incubated with 50 mg IgG-coated magnetic Dynabeads (Invitrogen) for 30 min. The beads were washed with lysis buffer twice and incubated with TEV protease in cleavage buffer (10 mM Tris-HCl, pH 8.0, 100 mM NaCl and 2 mM DTT) at 4 °C for 4–8 h. The released complex was concentrated to ~50 µl.

### Cryo-EM data collection and image processing

To prepare vitrified specimen, 3 µl of sample (OD_260_ = 3-5) was applied to glow-charged holey carbon grids (GiG R422) at 4 °C and 100% humidity using an FEI Vitrobot Mark IV. The grids were blotted for 1–3 s and rapidly plunged into liquid ethane. The cryogenic sample was first screened on a Talos F200C 200kV electron microscope equipped with a Ceta camera (FEI). A dataset was collected with a pixel size of 2.27 Å. Images were processed with RELION (50, 51). Particles were picked, cleaned and subjected to 3D classification using the mature yeast 40S ribosome as the initial model. A density map with clear structural feature was obtained at 11.7 Å resolution. This map was used as the initial model for subsequent image processing.

High resolution images were recorded on a Titan Krios (FEI) operated at 300 kV and equipped with a Falcon III detector. 984 images were collected with SerialEM (52) in the low-dose mode at a pixel size of 1.42 Å. Images were recorded in 23 movie frames with an exposure time of 1.5 s and a total dose of 30 e/ Å^2^ and corrected for motion with MOTIONCORR (53). Contrast transfer function (CTF) values were measured with CTFFIND4 (54). 925 good-quality images with a defocus range of 1–4 µm were chosen for particle picking in RELION. Particles were extracted with a box size of 360 pixels and downsized by 4 folds for 3D classification. Three rounds of 3D classification were performed with a mask of 400 Å diameter (Figure S2). The classes with high resolution and similar features were combined for further classification or refinement. The final 3D refinement was conducted on 98,155 particles with the gold-standard Fourier shell correlation (FSC) method (55). A density map at 3.73 Å resolution was obtained after correction of modulation transfer function of the detector and B-factor sharping. The reported resolution was based on the gold-standard FSC = 0.143 criterion. Local resolution was estimated with ResMap (56).

### Model building

The cryo-EM structure of Nog2-TAP pre-60S (PDB code 3JCT) was fit into the density as the starting model (18). The RNA and protein components without density were removed. The unmodeled density were assigned with assistance of the BALBES-MOLREP pipeline (57) and the CXMS data. The crystal structures of the Brx1 and Ebp2 complex determined here and Nsa1(45) were fitted as rigid body. Homology models of Rpf1 and Has1 were created by SWISS-MODEL (58). Rrp1 and Mak16 were manually built. A few unassigned densities were modeled as poly-alanine chain and additional weak densities were not modeled. Nucleotides 438–493 of helix ES7a were modeled based on prediction of RNAComposer (59). The structure was further adjusted in COOT (60) and refined in real space with secondary structure and geometry restraints using PHENIX (61). Structural figures were prepared with PyMOL 1.7 (Schrödinger, LLC) and Chimera (62).

### Expression and purification of the Brx1 and Ebp2 complex

The genes of Brx1 (total 291 residues) and Ebp2 (total 427 residues) were amplified from yeast genomic DNA and cloned with the transfer-PCR approach (Erijman et al., 2011). Ebp2 and its fragments were cloned into a modified pET28a vector (Novagen) and fused to an N-terminal His_6_-SMT3 tag. Brx1 and its fragments were cloned into a modified pETDuet-1 vector and fused to an N-terminal His_6_-GST tag followed by a PreScission cleavage site. The constructed plasmids were verified by DNA sequencing. Native or selenomethionine (SeMet)-labeled Brx1 and Ebp2 were co-expressed in *E. coli* Rosetta 2(DE3) strain and co-purified with HisTrap, heparin and gel filtration chromatography following the previously described procedure (63). The interaction between the two proteins was confirmed by GST pull-down assay.

### Crystallization and structural determination of the Brx1 and Ebp2 complex

Several fragments of Ebp2 and Brx1 were tested for crystallization. High quality crystals were grown from a complex containing residues 186–295 of Ebp2 and residues 26–259 of Ebp2. Both the native and SeMet-labeled complex (15 mg/ml in 10 mM Tris-HCl, pH 8.0 and 500 mM NaCl) were crystallized in 0.1 M Bis-Tris, pH 5.5 and 1.8 M ammonium sulfate with the vapor diffusion hanging drop method. The crystals were cryoprotected in 20% glycerol made in the reservoir solution and flash frozen in liquid nitrogen.

Diffraction data were collected at the beamline BL17U of Shanghai Synchrotron Radiation Facility and processed with HKL2000 (64). Two types of crystals that belonged to the C2_1_ or P2_1_ space group were obtained at the same condition. The structure was first solved by the single-wavelength anomalous diffraction (SAD) method based on a C2_1_ Se-derivative dataset at 2.8 Å resolution using SHARP (65) (Table S3). The resultant electron density map was of moderate quality. The experimental phases were iteratively combined with the phases calculated from partially built model to improve the map. A better dataset at 2.3 Å resolution was later collected on the native P2_1_ crystal and used for final refinement. The model was built in COOT(60) and refined with PHENIX(61). The final model contains two copies of the complex composed of Ebp2 residues 195–293 and Brx1 residues 26–65, 70–195 and 210–255, 308 water molecules and one sulfate ion.

### Mass spectrometric analysis

Mass spectrometric analysis was conducted as previously described (2). The total spectral counts per 100 residues (SCPHR) were calculated for each identified protein and further normalized against Brx1, Ebp2, Erb1, Ytm1, Nop7, Cic1 and Has1, yielding the relative spectral abundance factor (RSAF) (Supplementary Dataset 1).

### CXMS

CXMS was performed as described (7). The Rpf1-TAP sample containing ~10 μg total proteins was crosslinked with BS^3^ or DSS, precipitated with acetone and digested by trypsin. The sample was analyzed by LC-MS/MS on an EASY-nLC 1000 system interfaced to a Q-Exactive HF mass spectrometer (Thermo Fisher Scientific). Cross-linked peptides were identified with pLink (Yang et al. 2012).

### Protein interaction analysis of Rpf1 and Mak16

Yeast two-hybrid and GST pull-down assays were conducted as described (66). Rpf1 was cloned into a modified pET28a vector (Novagen) and fused to an N-terminal His_6_-SMT3 tag. Mak16 and its fragments were cloned into a modified pETDuet-1 vector and fused to an N-terminal His_6_-GST tag followed by a PreScission cleavage site. For GST pull-down assay, Rpf1 was expressed and purified by HisTrap chromatography, Ulp1 cleavage of the His_6_-SMT3 tag, heparin and gel filtration chromatography. Mak16 was purified by HisTrap chromatography without cleavage of the tag. To purify the complex, Rpf1 and Mak19 were co-expressed and co-purified following the previously described procedure (63).

### Accession Number

The cryo-EM density map, coordinates and structural factors have been deposited to EMDB and PDB under accession numbers EMD-XXXX, XXXX and 5Z1G.

## Acknowledgments

We thank the Center for Biological Imaging (CBI), Institute of Biophysics, Chinese Academy of Science for cryo-EM study and HPC-Service Station in CBI for image processing. We are grateful to Fei Sun, Xiaojun Huang, Zhenxi Guo, Gang Ji, Bolin Zhu, Shuoguo Li and Deyin Fan at CBI for help in EM sample preparation and data collection, Gaihong Cai and She Chen at the Proteomics Center of NIBS for mass spectrometry analysis and the staffs at beamline BL17U of National Facility for Protein Sciences Shanghai and Shanghai Synchrotron Radiation Facility for assistance in data collection. The study was supported by Strategic Priority Research Program of Chinese Academy of Sciences [XDB08010203], National Key R&D Program of China [2017YFA0504600], National Natural Science Foundation of China [31430024, 91540201, 31325007], the Ministry of Science and Technology of China [973 grant 2014CB84980001 to M.-Q.D.] and 100 Talents Program of CAS.

---

**Figure S1.**
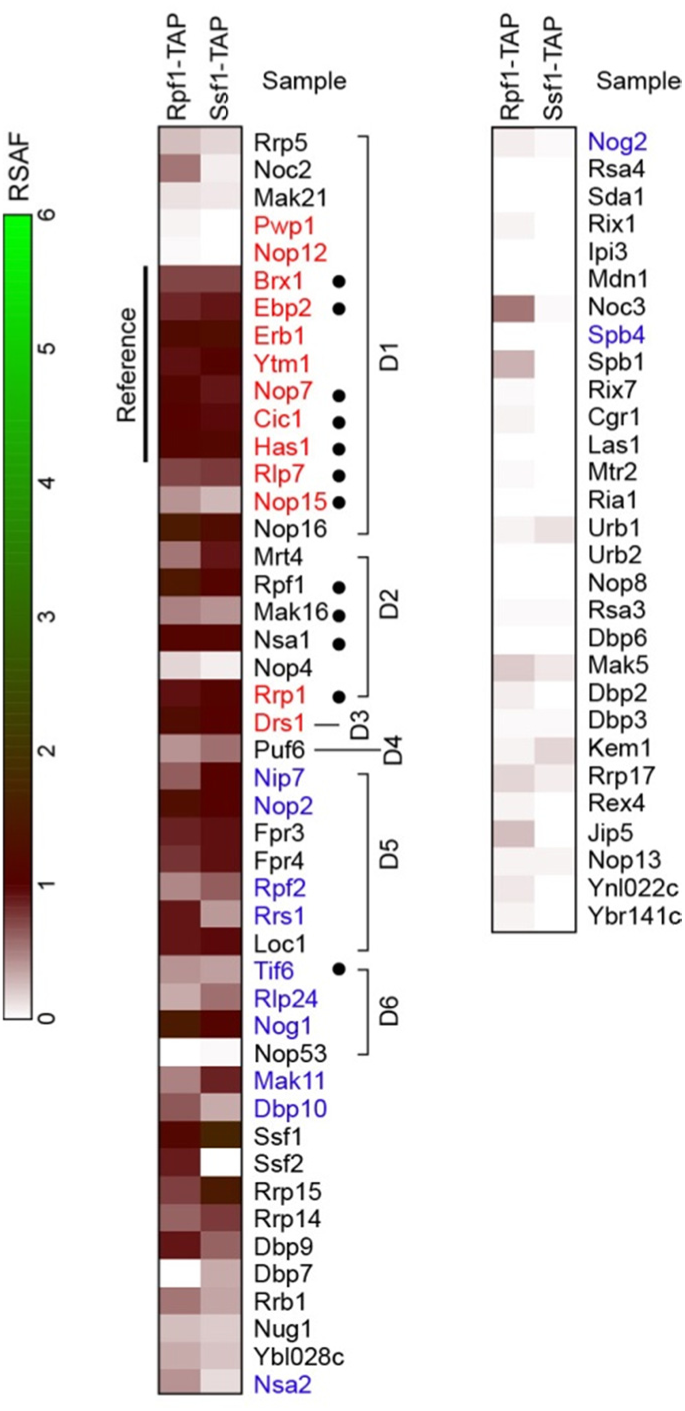
Heatmap of pre-60S AFs in Rpfl-TAP particle. The proteins identified by mass spectrometry are color coded according to their RSAF values normalized against the reference proteins Brx1, Ebp2, Erb1, Ytm1, Nop7, Cic1 and Has1. The previously reported Ssf1-TAP data were included for comparison (14). The AFs on the top of the list are arranged by their association order to pre-27S rRNA fragments ending at domains I to VI (D1-D6). A3-factors and B-factors are colored red and blue, respectively. Solid circles mark 12 AFs modeled in the Rpf1-TAP pre-60S structure.

**Figure S2.**
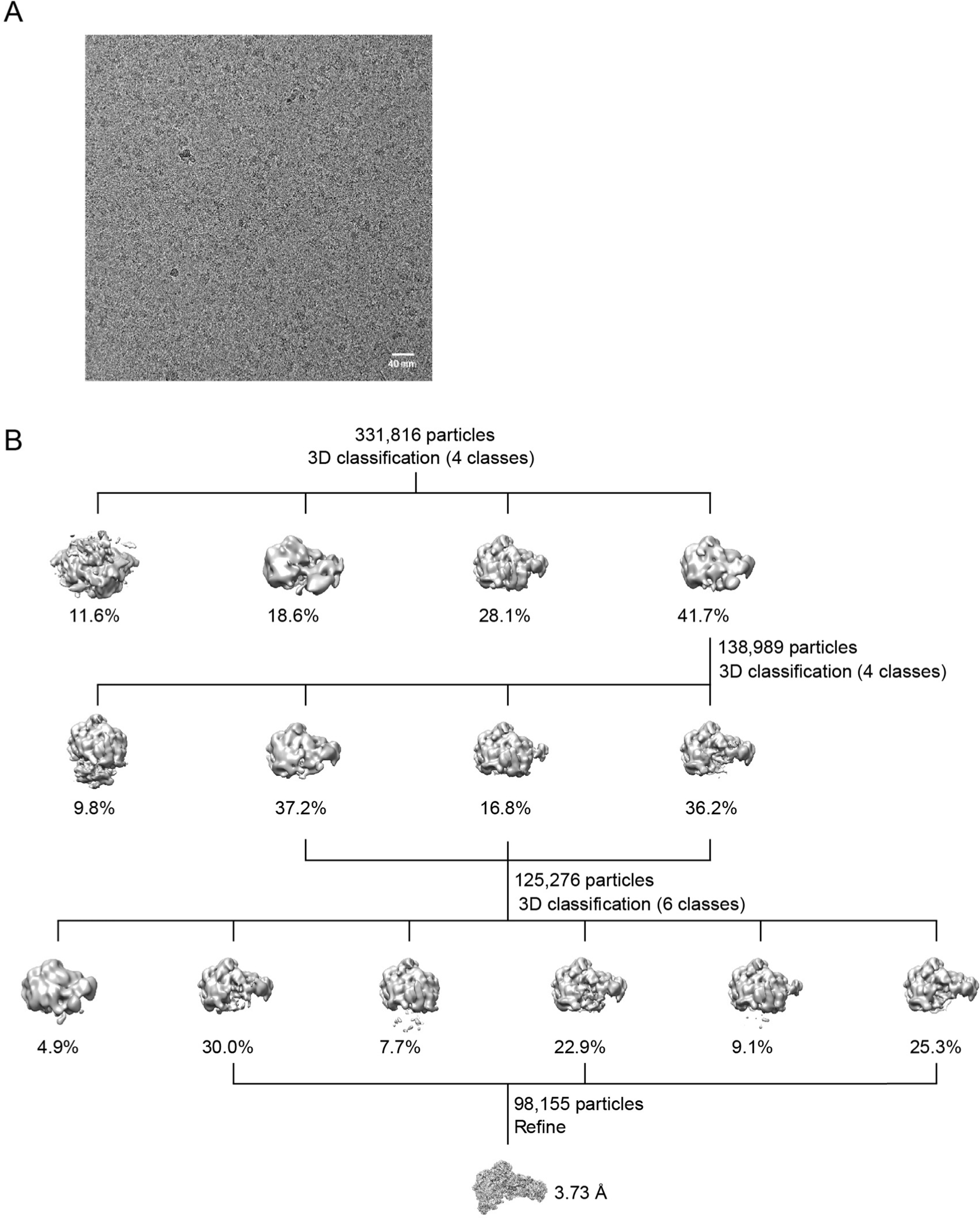
Cryo-EM analysis of Rpf1-TAP particles. (A) Electron micrograph of Rpf1-TAP particles. Bar = 40 nm. (B) Flowchart of 3D classification and refinement.

**Figure S3.**
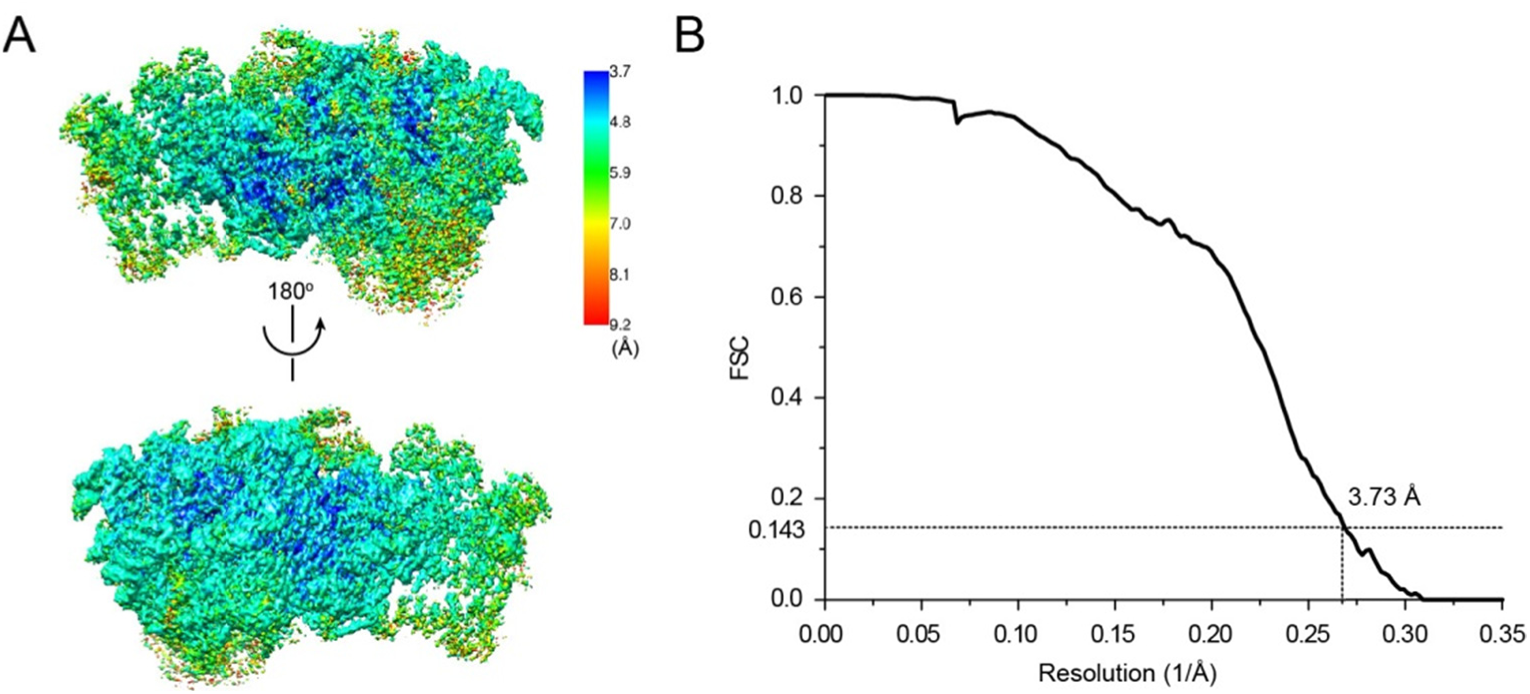
Resolution of the cryo-EM map of Rpf1-TAP pre-60S. (A) Two opposite views of local resolution cryo-EM map. (B) Gold-standard Fourier shell correlation curve for the construction.

**Figure S4.**
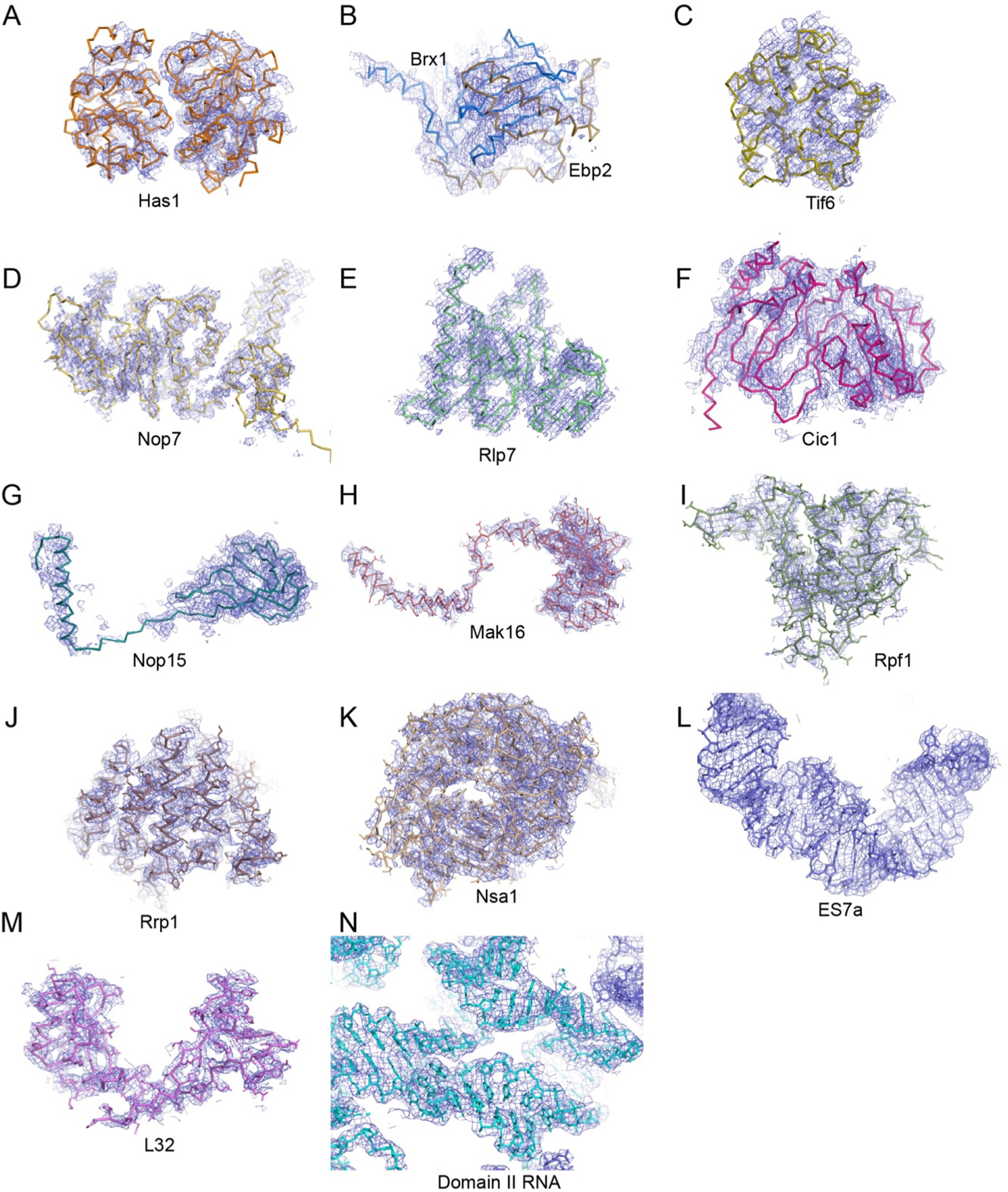
Representative cryo-EM densities with fitted structural models. (A) Has1. (B) The Brx1-Ebp2 complex. (C) Tif6. (D) Nop7. (E) Rlp7. (F) Cic1. (G) Nop15. (H) Mak16. (I) Rpf1. (J) Rrp1. (K) Nsa1. (L) ES7a. (M) L32. (N) Domain II RNA. To reduce noise, the Gaussian filtered map (standard deviation =1) was displayed for Has1, Brx1/Ebp2 and Tif6. Side chains of protein residues were shown for Rpf1, Mak16, Rrp1, Nsa1 and L32.

**Figure S5.**
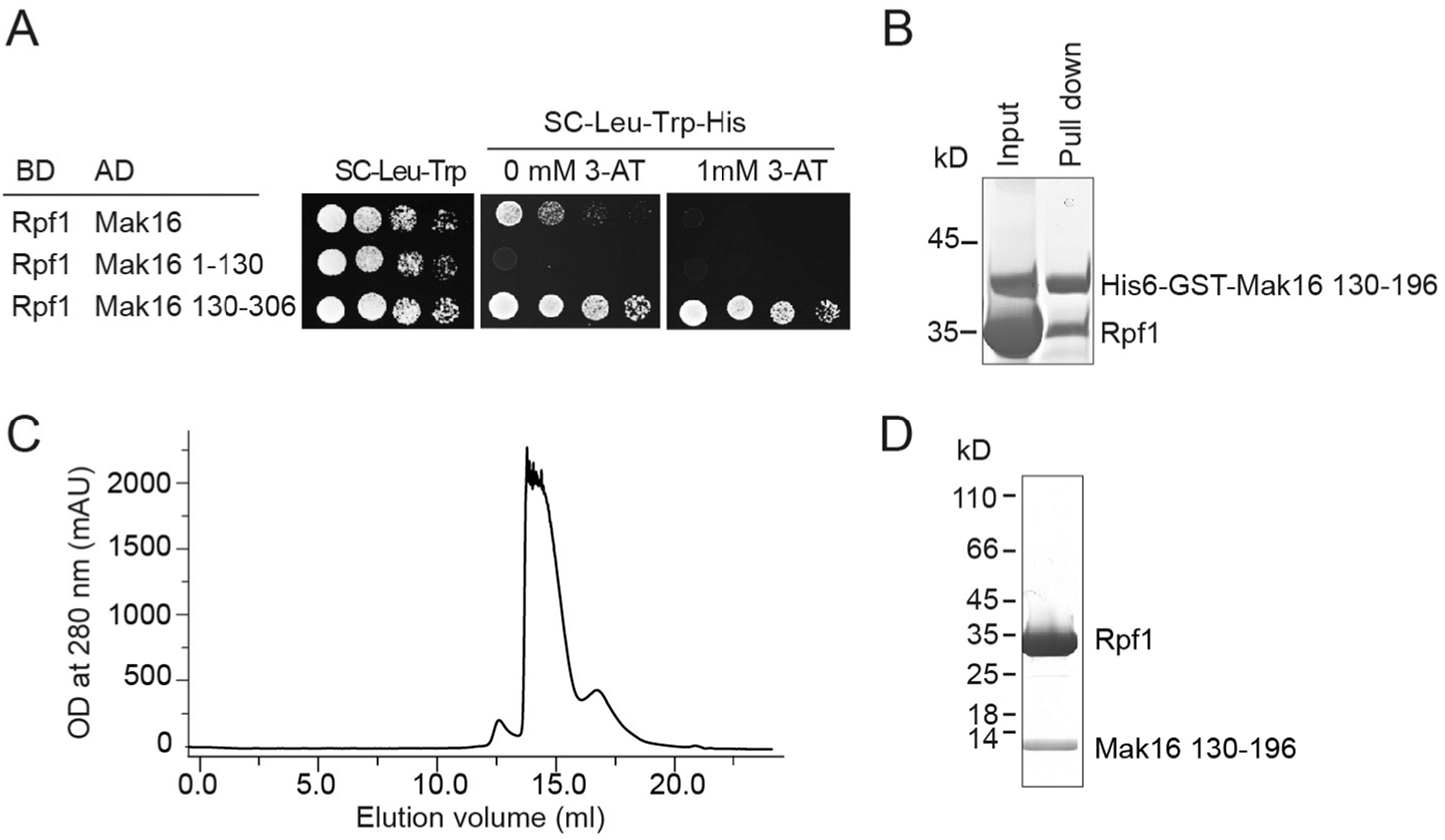
Rpfl binds a short sequence of Mak16. (A) Yeast two hybrid assay. Rpfl was fused to the GAL4 DNA-binding domain (BD) as bait. Mak16 and its fragments were fused to the GAL4 activation domain (AD) as prey. Yeast AH109 cells were co-transformed with bait and prey plasmids, five-fold serially diluted and spotted in plates with Synthetic Complete (SC) medium lacking Leu and Trp as growth controls and in plates with SC medium lacking Leu, Trp and His and containing the indicated concentration of 3’-amino-1,2,4-triazole (3-AT) to examine the prey-bait interaction. (B) GST pull-down assay. Rpf1 was incubated with a His_6_-GST-tagged Mak16 fragment containing residues 130–196 and pulled down with glutathione Sepharose beads. The input and eluate were examined by SDS-PAGE and Coomassie blue staining. The positions of molecular standards are indicated on the left. (C) Gel filtration profile of co-expressed and co-purified complex of Rpf1 and Mak16 130–196 in a Superdex 200 column. (D) SDS-PAGE gel of the peak fraction in C.

**Table S1.**
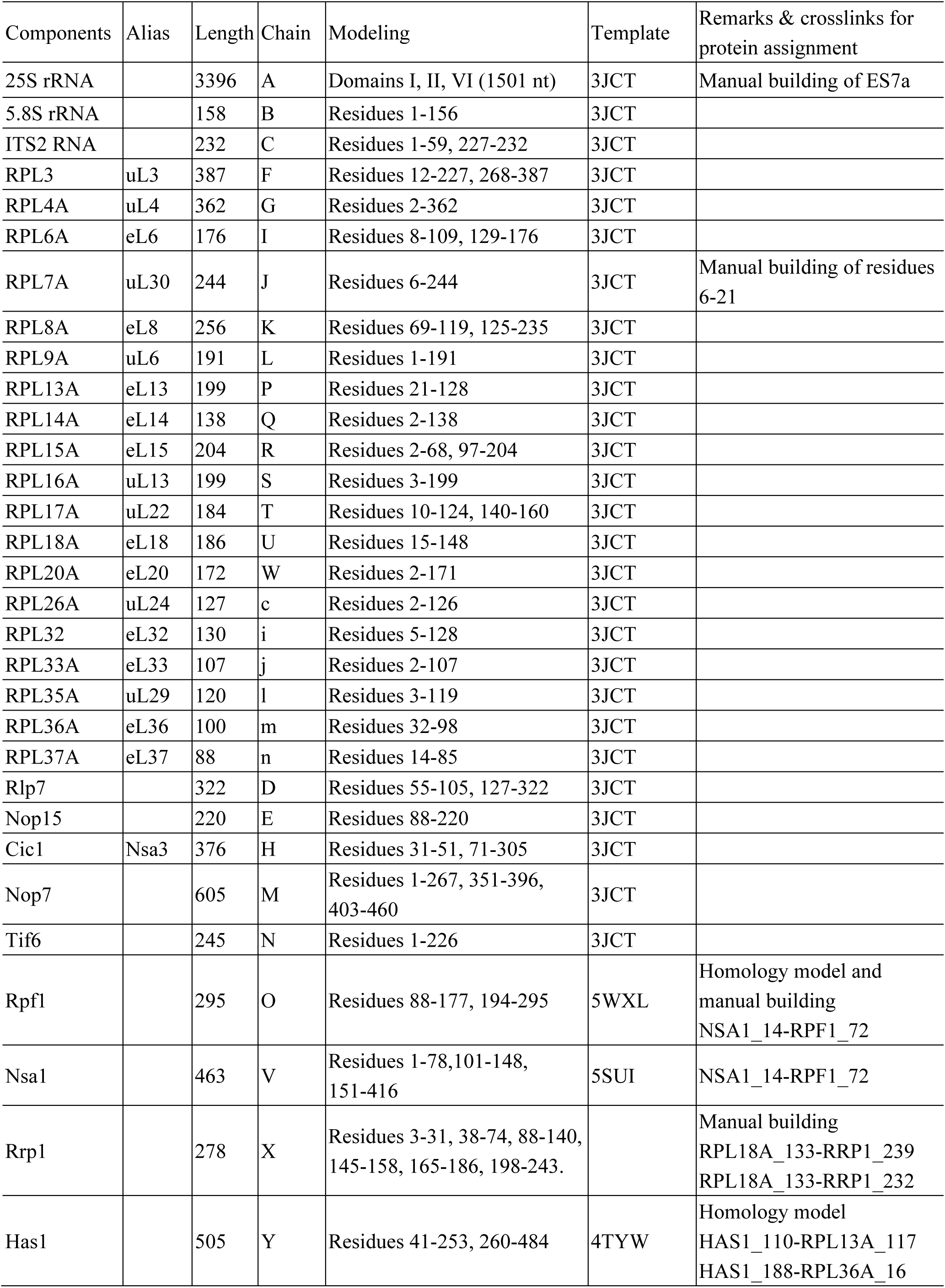
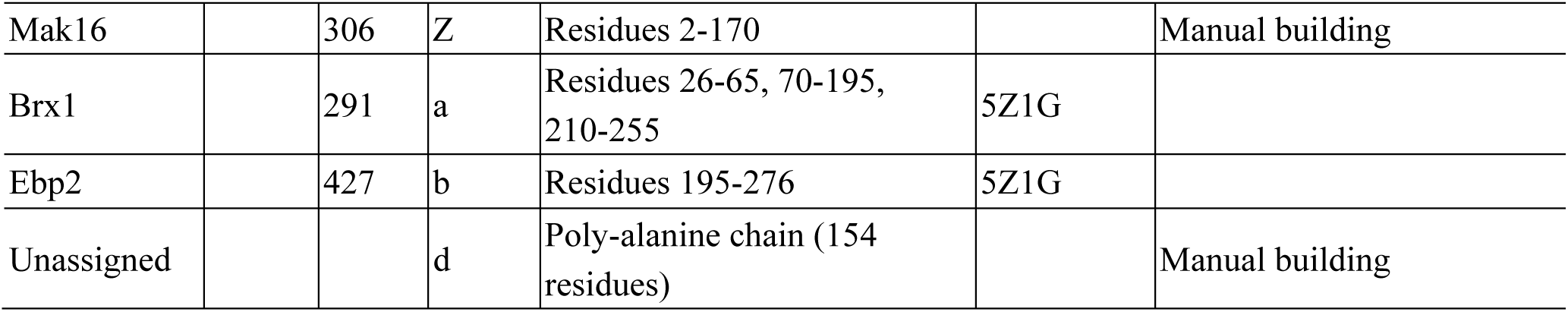
Modeling of the Rpf1-TAP pre-60S structure

| Components | Alias | Length | Chain | Modeling | Template | Remarks & crosslinks for protein assignment |
| --- | --- | --- | --- | --- | --- | --- |
| 25S rRNA |  | 3396 | A | Domains I, II, VI (1501 nt) | 3JCT | Manual building of ES7a |
| 5.8S rRNA |  | 158 | B | Residues 1-156 | 3JCT |  |
| ITS2 RNA |  | 232 | C | Residues 1-59, 227-232 | 3JCT |  |
| RPL3 | uL3 | 387 | F | Residues 12-227, 268-387 | 3JCT |  |
| RPL4A | uL4 | 362 | G | Residues 2-362 | 3JCT |  |
| RPL6A | eL6 | 176 | I | Residues 8-109, 129-176 | 3JCT |  |
| RPL7A | uL30 | 244 | J | Residues 6-244 | 3JCT | Manual building of residues 6-21 |
| RPL8A | eL8 | 256 | K | Residues 69-119, 125-235 | 3JCT |  |
| RPL9A | uL6 | 191 | L | Residues 1-191 | 3JCT |  |
| RPL13A | eL13 | 199 | P | Residues 21-128 | 3JCT |  |
| RPL14A | eL14 | 138 | Q | Residues 2-138 | 3JCT |  |
| RPL15A | eL15 | 204 | R | Residues 2-68, 97-204 | 3JCT |  |
| RPL16A | uL13 | 199 | S | Residues 3-199 | 3JCT |  |
| RPL17A | uL22 | 184 | T | Residues 10-124, 140-160 | 3JCT |  |
| RPL18A | eL18 | 186 | U | Residues 15-148 | 3JCT |  |
| RPL20A | eL20 | 172 | W | Residues 2-171 | 3JCT |  |
| RPL26A | uL24 | 127 | c | Residues 2-126 | 3JCT |  |
| RPL32 | eL32 | 130 | i | Residues 5-128 | 3JCT |  |
| RPL33A | eL33 | 107 | j | Residues 2-107 | 3JCT |  |
| RPL35A | uL29 | 120 | l | Residues 3-119 | 3JCT |  |
| RPL36A | eL36 | 100 | m | Residues 32-98 | 3JCT |  |
| RPL37A | eL37 | 88 | n | Residues 14-85 | 3JCT |  |
| Rlp7 |  | 322 | D | Residues 55-105, 127-322 | 3JCT |  |
| Nop15 |  | 220 | E | Residues 88-220 | 3JCT |  |
| Cic1 | Nsa3 | 376 | H | Residues 31-51, 71-305 | 3JCT |  |
| Nop7 |  | 605 | M | Residues 1-267, 351-396, 403-460 | 3JCT |  |
| Tif6 |  | 245 | N | Residues 1-226 | 3JCT |  |
| Rpf1 |  | 295 | O | Residues 88-177, 194-295 | 5WXL | Homology model and manual building<br>NSA1_14-RPF1_72 |
| Nsa1 |  | 463 | V | Residues 1-78,101-148, 151-416 | 5SUI | NSA1_14-RPF1_72 |
| Rrp1 |  | 278 | X | Residues 3-31, 38-74, 88-140, 145-158, 165-186, 198-243. |  | Manual building<br>RPL18A_133-RRP1_239<br>RPL18A_133-RRP1_232 |
| Has1 |  | 505 | Y | Residues 41-253, 260-484 | 4TYW | Homology model<br>HAS1_110-RPL13A_117<br>HAS1_188-RPL36A_16 |
| Mak16 |  | 306 | Z | Residues 2-170 |  | Manual building |
| Brx1 |  | 291 | a | Residues 26-65, 70-195,<br>210-255 | 5Z1G |  |
| Ebp2 |  | 427 | b | Residues 195-276 | 5Z1G |  |
| Unassigned |  |  | d | Poly-alanine chain (154<br>residues) |  | Manual building |

**Table S2.**
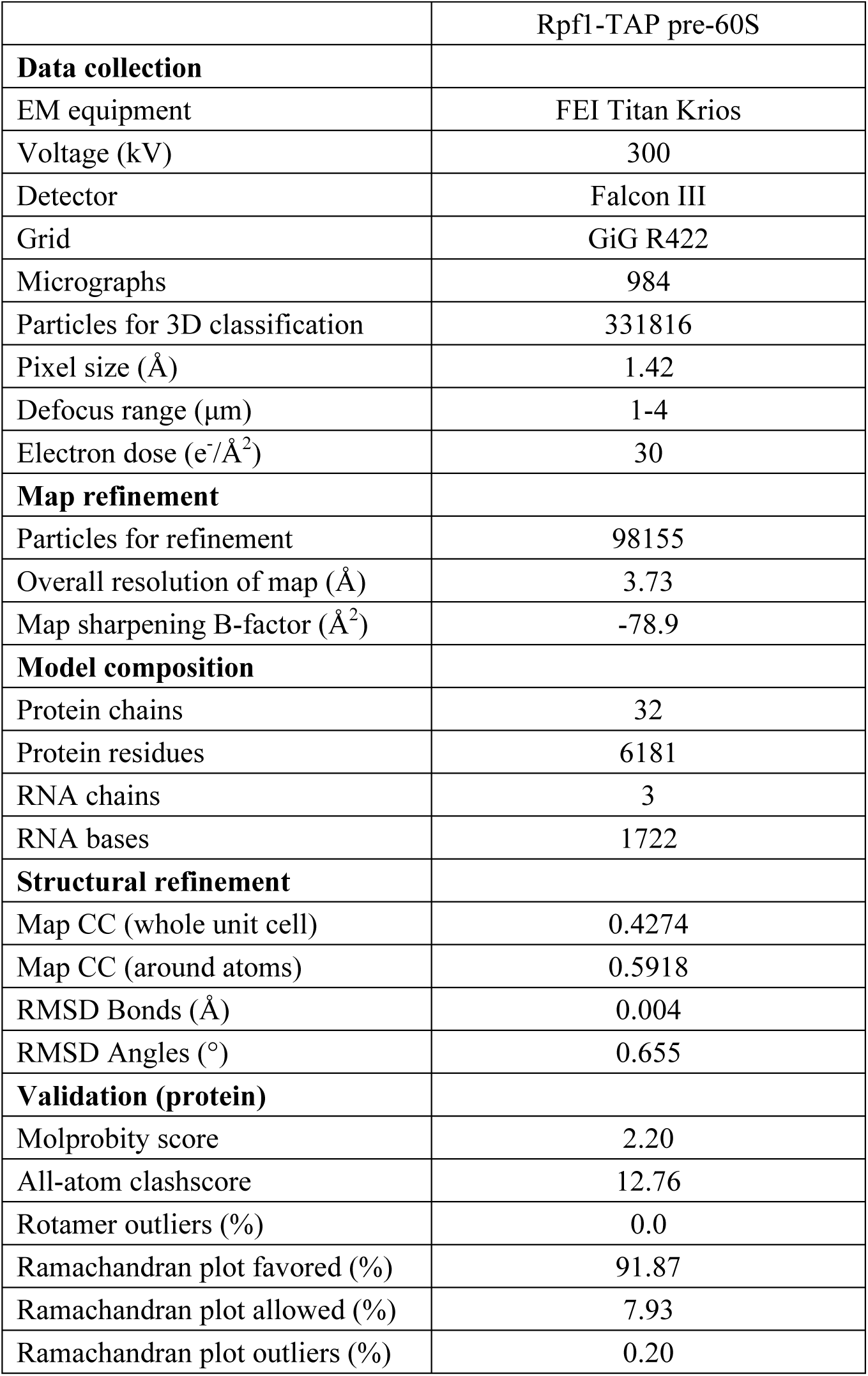
Statistics of data collection, structural refinement and model validation

**Table S3.** Data collection and refinement statistics of the Brx-Ebp2 crystal structure

| Crystal form | Native | Se-labeled |
| --- | --- | --- |
| Data collection |  |  |
| Space group | P2<sub>1</sub> | C2<sub>1</sub> |
| Cell dimensions |  |  |
| a, b, c (Å) | 80.9, 47.4, 100.7 | 100.3, 47.0, 71.9 |
| $\alpha$, $\beta$, $\gamma$ (°) | 90.0, 94.0, 90.0 | 90.0, 118.8, 90.0 |
| Wavelength (Å) | 0.97930 | 0.97930 |
| Resolution range (Å) | 30.0-2.3(2.34-2.30) | 25.0-2.80(2.85-2.80) |
| Unique reflections | 33644(1715) | 7463 (348) |
| Redundancy | 3.5(3.6) | 5.9(6.2) |
| $\langle I \rangle / \langle \sigma(I) \rangle$ | 17.8(4.0) | 20.3(4.5) |
| Completeness (%) | 97.5(99.9) | 99.7(99.4) |
| $R_{\text{merge}}$ | 0.099(0.382) | 0.179(0.784) |
| Structure refinement |  |  |
| Resolution range (Å) | 25-2.3(2.36-2.3) |  |
| No. reflections | 33214 |  |
| No. atoms | 5459 |  |
| Protein | 5146 |  |
| Water | 308 |  |
| Sulfate ion | 5 |  |
| $R_{\text{work}}$ | 0.230(0.278) | |
| $R_{\text{free}}$ | 0.297(0.394) | |
| Mean B factor (Å<sup>2</sup>) | 25.96 |  |
| Rmsd bond length (Å) | 1.091 |  |
| Rmsd bond angles (°) | 0.008 |  |
| Ramachandran Plot |  |  |
| Favored (%) | 96.53 |  |
| Allowed (%) | 3.14 |  |
| Outliers (%) | 0.33 |  |

Values in parentheses are for the data in the highest resolution shell.

## Supplementary Dataset 1. Mass spectrometry data

The SCPHR and RSAF values of identified proteins are displayed in two separate sheets. The RSAF is normalized against Brx1, Ebp2, Erb1, Ytm1, Nop7, Cic1 and Has1. The top rows include the summed SCPHR or RSAF values for 90S proteins, pre-40S proteins, pre-60S proteins, small subunit ribosomal proteins (RPS), large subunit ribosomal proteins (RPL) and total identified proteins, the total spectral counts (SpC) of reference proteins and the molar percentages of pre-60S AFs and RPLs over all detected proteins. (Excel table)

## Supplementary Dataset 2. CXMS data of Rpf1-TAP sample

The crosslinked proteins, crosslinked peptides, spectral counts and best E-values are listed. (Excel table)

